# Neoteny in visual system development of the spotted unicornfish, *Naso brevirostris* (Acanthuridae)

**DOI:** 10.1101/691774

**Authors:** Valerio Tettamanti, Fanny de Busserolles, David Lecchini, Justin Marshall, Fabio Cortesi

## Abstract

Ontogenetic changes of the visual system are often correlated to shifts in habitat and feeding behaviour of animals. Coral reef fishes begin their lives in the pelagic zone and then migrate to the reef. This transition of habitat frequently involves a change in diet and light environment as well as major morphological modifications. The spotted unicornfish, *Naso brevirostris*, is known to shift diet from zooplankton to algae and back to zooplankton when transitioning from larval to juvenile and then to adult stages. Concurrently, *N. brevirostris* also moves from an open pelagic to a coral-associated habitat before migrating up in the water column when reaching adulthood. Using retinal mapping techniques, we discovered that the distribution and density of ganglion and photoreceptor cells in *N. brevirostris* do not change with the habitat or the feeding habits of each developmental stage. Instead, fishes showed a neotenic development with a slight change from larval to juvenile stages and not many modifications thereafter. Visual gene expression based on RNA sequencing mirrored this pattern; independent of stage, fishes mainly expressed three cone opsin genes (*SWS2B, RH2B, RH2A*), with a quantitative difference in the expression of the green opsin genes (*RH2A* and *RH2B*) when transitioning from larvae to juveniles. Hence, contrary to the ontogenetic changes found in many animals, the visual system is fixed early on in *N. brevirostris* development calling for a thorough analysis of visual system development of the reef fish community.

## Introduction

Many animals use vision to perform important behavioural tasks such as feeding, mating, avoiding predators and to find a suitable home (Cronin *et al.* 2014). At the core of the vertebrate visual system is the retina, an extrusion of the brain which is subdivided into various functional layers, two of which are at the centre of this study, the photoreceptor layer and the ganglion cell layer.

The photoreceptor layer is the first stage of visual processing and is composed of morphologically diverse cone and rod photoreceptor cells which absorb light, transform it into an electrical signal, and send the information downstream to various neural cells via the phototransduction cascade. Cones mediate vision in bright light conditions and colour vision while rods mediate vision in dim light conditions (Walls 1942). Cones can further be classified into different types depending on their morphology and/or the type of photopigment (an opsin protein covalently bound to a light absorbing chromophore) they possess (Hunt *et al.* 2014). Morphologically, cones can be classified as single, double, triple or quadruple, although only the first two configurations are common and are often arranged in regular and specific patterns or mosaics (Peichl *et al.* 2004; Bowmaker and Kunz 1987). Molecularly, cones are also classified into four types based on the opsin genes they express that encode for different protein classes sensitive to different parts of the visible light spectrum. The short-wavelength protein class 1 opsin (SWS1) maximally sensitive to UV-violet wavelengths (355-450 nm λ_max_), and a second short-wavelength class opsin (SWS2) maximally sensitive to the violet-blue part of the spectrum (410-490 nm λ_max_), are expressed in single cones. Double cones express middle-wavelength class 2 opsin (RH2) maximally sensitive to blue-green wavelengths (470-535 nm λ_max_), and a long-wavelength class opsin (LWS) maximally sensitive to the green-red part of the light spectrum (490-570 nm λ_max_). Most vertebrates possess a single type of rod photoreceptor expressing the rod opsin protein (RH1; 460-530 nm λ_max_) (Bowmaker 2008; Walls 1934).

The ganglion cell layer is the last stage of visual processing in the retina and is composed of ganglion cells that possess axons that reach to the inner surface of the retina and converge into the optic nerve to send the information into the central nervous system (Walls 1942). Therefore, the arrangement and the spacing between one ganglion cell to another is one of the determining factors of visual acuity (or resolution) (Fernald 1988).

In order to perform at its optimum, the visual system of a particular species is adapted to the type of habitat they live in and to the prevailing surrounding light conditions (Lythgoe 1979). In general, vertebrates range from cone-monochromats with a single spectral class of cone photoreceptor (e.g., sharks and many rays), over di- and trichromats (e.g., most mammals and many marine fishes), to tetrachromats (e.g., most birds and many freshwater fishes; Bowmaker 2008). Cone photoreceptors and their respective opsin repertoires are particularly diverse is teleost fishes (e.g., Musilova *et al.* 2019; Lin *et al.* 2017). This is thought to primarily be due to the different light environments fishes inhabit (Lythgoe 1979; Cronin *et al.* 2014), but in some instances may also be driven by sexual selection (Endler 1990) and/or the feeding habits of species. For example, UV photoreception increases feeding efficiency in some fishes eating UV-absorbing or scattering zooplankton (Loew *et al.* 1993; Novales-Flamarique and Hawryshyn 1994; Flamarique 2016), while herbivorous fishes may profit from visual systems tuned to longer wavelengths due to the red-reflecting properties of chlorophyll (Marshall *et al.* 2003; Stieb *et al.* 2017; Cortesi *et al.* 2018).

Further to the type, the density of photoreceptors and ganglion cells can also vary not only between species, but also within an individual’s retina (Shand *et al.* 1999; Shand *et al.* 2000). The study of their distribution using the wholemount technique (Stone and Johnston 1981; Ullmann *et al.* 2012; Coimbra *et al.* 2006) provides useful information on the visual ecology of a species, which usually reflects its habitat and behavioural ecology (Hughes *et al.* 1977; Bozzano and Collin 2000; Collin and Pettigrew 1988b, 1988a; Collin and Pettigrew 1989). Two main specializations can be found in vertebrates: area and streaks, (Collin and Pettigrew 1988b, 1988a). Both specializations have higher densities of cells compared to the rest of the retina, resulting in regions of acute vision in the corresponding field of view. An area is a concentric increase in cell density in a particular region of the retina, in some vertebrates it is termed a fovea due to other structural adaptations (Walls, 1942). In teleost fishes, areas are often located temporally (i.e. area temporalis) and found in species that live in enclosed environments such as caves or coral structures, and/or coral overhangs (Collin and Shand 2003; Collin and Pettigrew 1989). The temporal area receives the visual information from the frontal field of view, corresponding to the natural swimming direction of fishes. Nevertheless, multiple area centralis can also be found in a single retina (Collin and Pettigrew 1989). For example, Triggerfishes (Balistidae) possess an area in both the nasal and temporal part of the retina, which correlates with two main visual tasks: feeding (temporal) and predator avoidance (nasal) (Ito and Murakami 1984; Collin and Pettigrew 1988b).

A horizontal streak is defined by an increase in cell density along the meridian. Most horizontal streaks are found in the central meridian, but sometimes they can also be located more ventrally or dorsally (Collin and Shand 2003). The streak maintains a high spatial resolving power throughout the horizontal section of the retina and is thought to be used to scan the horizon. It leads to an elongated sampling of the visual environment without continuous eye movements (Collin and Shand 2003). Teleost fishes possessing a horizontal streak are commonly found in open water environments such as sandy bottoms or pelagic open ocean environments (Collin and Pettigrew 1988b).

Variability in retinal structure and opsin gene repertoire does not only exist between species but both visual features may also change throughout the life of an individual. This is especially true for species that undergo substantial habitat changes during ontogeny such as coral reef fishes. The life of most coral reef fishes starts in the shallower zone of the open ocean as larvae (Helfman *et al.* 2009; Job and Bellwood 2000), where resources may be high and the risk of predation is low (Fortier and Harris 1989). At this stage, pelagic fish larvae feed typically on zooplankton (Boehlert 1996) and rely on vision primarily for fundamental tasks such as predator avoidance and feeding (Leis and Carson-Ewart 1999). After their oceanic phase, reef fish larvae typically find a suitable coral reef patch to settle on and again vision is one of the main senses used (Lecchini *et al.* 2005a; Lecchini *et al.* 2005b). During this settlement phase, reef fish larvae undergo metamorphosis and reach the juvenile stage in which they usually already possess all basic morphological features of the adult form (Holzer *et al.* 2017). Following settlement on the reef, juvenile fishes are challenged with visual cues that are much more complex than in the open ocean varying both in chromaticity and luminance. Hence, at this stage (or slightly before – Cortesi *et al.* 2016) the visual system of coral reef fishes is expected to undergo changes both in morphology and physiology (Helfman *et al.* 2009).

To date, changes in arrangement (i.e. mosaic) and distribution of the photoreceptor cells throughout ontogeny have been documented only in few coral reef fishes (Shand 1997). These changes are thought to enhance survivability by increasing feeding success and facilitate predator avoidance in reef stages (juveniles and adults; Shand 1997). Along with changes in morphology, ontogenetic changes in opsin gene expression have also been reported from a handful of species (Cortesi *et al.* 2016; Cortesi *et al.* 2015b). For example, in the dottyback *Pseudochromis fuscus*, the number and type of opsin genes that are expressed differs between larval, juvenile, and adult stages. Along with the change in opsin gene expression, the visual system may also transform to more complex colour processing capability, such as di-to tri-chromacy or even up to tetrachromacy. This increase in chromaticity ultimately requires behavioural testing to confirm and is likely to reflect major habitat transitions throughout development equipping e.g., dottybacks with a more complex visual system as they grow and mature (Cortesi *et al.* 2016; Cortesi *et al.* 2015b). In comparison, while some freshwater cichlid species show a similarly dynamic change in opsin gene expression through ontogeny, other species do not change gene expression much (neoteny) or then, they directly develop from the larval to the adult gene-expression pattern (Carleton *et al.* 2008; Härer *et al.* 2017). We currently do not know whether a progressive developmental change of the visual system, as e.g., found in the dottyback (Cortesi *et al.* 2016), is a common feature shared among reef fishes, or whether some species also show different developmental modes, similar to what is found in cichlid fishes.

In this study we investigated ontogenetic changes in retinal topography and opsin gene expression in three life stages (larval, juvenile, adult) of the spotted unicornfish, *Naso brevirostris*, from the surgeonfish family (Acanthuridae) (Fig. 1). *N. brevirostris* is known to shift both diet and habitat during ontogeny (Choat *et al.* 2002; K. Clements, D. Bellwood personal communication). Pelagic larvae feed on zooplankton before settling on the reef where they mainly feed on algae as juveniles. As adults, *N. brevirostris* migrate to the reef slope returning to a zooplanktivorous diet (Choat *et al.* 2002; Choat *et al.* 2004). We therefore hypothesized that the visual system of *N. brevirostris* would show a ‘classic’ developmental mode, linked to either changes in habitat or diet, or both, and with a progression from larval, to juvenile and finally adult traits. Moreover, *N. brevirostris* develops an elongated rostral snout during maturation, and this prominent morphological change may also affect its visual requirements as it might obstruct the visual field of the fish.

**Fig. 1.**
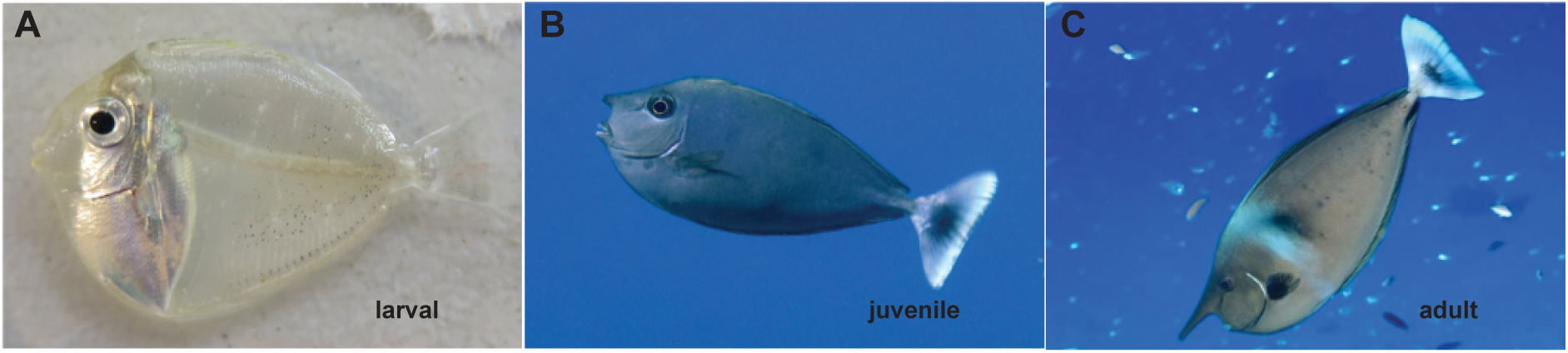
*Naso brevirostris* developmental stages. The spotted unicornfish, *N. brevirostris*, shows pronounced ontogenetic changes in habitat, diet and morphology. (A) A ‘transparent’ zooplanktivorous larva at the settlement stage (i.e., when returning from the pelagic to the reef). (B) An algivorous juvenile that lives in close proximity to the reef. (C) A zooplanktivorous adult that lives in the water column above the reef. Note the growth of the prominent snout throughout development.

## Materials and Methods

### Study species and collection

Individuals of *N. brevirostris* were collected on the Northern Great Barrier Reef, Australia, under the Great Barrier Reef Marine Park Association (GBRMPA) permits G17/38160.1 and G16/38497.1, Queensland Fisheries permit #180731, or in French Polynesia. Adults (n = 6) were collected with a spear gun from No Name Reef (14°65 ′S, 145°65′E) on the outer Great Barrier Reef, Australia, in February 2018. Juveniles (n = 8) were collected using barrier nets, spear guns or clove oil and hand nets from reefs surrounding Lizard Island (14°40′S, 145°27′E) on the Great Barrier Reef between February 2016 – February 2018. Two additional juvenile samples were acquired through the aquarium supplier Cairns Marine (http://www.cairnsmarine.com/). Larval fish (n = 5) were captured using a crest net on Tetiaroa Island, French Polynesia (16°99′S, 149°58′W) in March 2018 (Lecchini *et al.* 2004, Besson *et al.* 2017). All animals were quickly anaesthetised following the NHMRC Australian Code of Practice under an animal ethics protocol of The Queensland Brain Institute (QBI/236/13/ARC/US AIRFORCE and QBI/304/16).

Each individual was photographed with a ruler in the frame, to be able to extract the standard length later on using Fiji v.1.0 (Schindelin *et al.* 2012). The eyes were enucleated and the cornea and lens removed using micro-dissection scissors. A small dorsal cut was made to keep track of the eye’s orientation. The samples collected for retinal mapping were fixed in 4% paraformaldehyde in 0.1 M phosphate buffer (PBS; pH 7.4) and stored at 4 degrees Celsius and the eyes used for RNA sequencing were kept in RNAlater (Sigma) and stored at −20 degrees Celsius. For each eye, the lens diameter was measured after dissection and fixation.

### Preparation of retinal wholemounts

Retinal wholemounts were prepared according to standard protocols (Stone and Johnston 1981; Coimbra *et al.* 2006; Coimbra *et al.* 2012). Briefly, each eye cup was cut radially multiple times, to flatten it on a microscopy glass slide without damaging the tissue. The retina was oriented using the falciform process that extends ventrally. The sclera and choroid were gently removed and the retinas where bleached overnight in the dark at ambient temperature in a 3% hydrogen peroxide solution (in PBS). Large-sized adult retinas, that have a more developed retinal pigment epithelium, were bleached in the same solution but with a few drops of potassium hydroxide (Ullmann *et al.* 2012). While potassium hydroxide accelerates the bleaching process by increasing the pH of the solution, this type of bleaching is quite aggressive for the tissue. Therefore, these retinas were only bleached for 2-3h in the dark.

For ganglion cell analyses, retinas were mounted ganglion cell layer facing up on a gelatinized slide and left to dry overnight at room temperature in formalin vapours (Coimbra *et al.* 2006; Coimbra *et al.* 2012). Wholemounts were then stained in 0.1% cresyl violet (Nissl staining) following the protocol of Coimbra *et al.* (2006) and then mounted with Entellan New (Merck). Shrinkage of the retina using this technique is usually dimmed negligible and, if present, restricted to the borders of the retina (Coimbra *et al.* 2006). In this study however, all the retinas were not equal in shrinkage due to major differences in retina size between developmental stages. As such, the smaller retinas (larval stage) were more affected by shrinkage, due to their smaller surface (i.e. higher proportion of retinal borders), than the other stages (adult and juvenile stages). Shrinkage in these retinas was easily identified under the microscope and was taken into consideration in the data interpretation.

For photoreceptor analyses, retinas were wholemounted in 100% glycerol, on non-gelatinized slides with the inner (vitreal) surface facing downwards. Contrary to ganglion cell mounting, photoreceptor mounting shows negligible shrinkage as it takes place in an aqueous medium (Peichl *et al.* 2004).

### Stereological analyses and construction of topographic maps

The topographic distribution and the total number of ganglion cells, single cones, double cones and total cones in the three life stages of *N. brevirostris* were assessed using the optical fractionator technique (West *et al.* 1991), modified for wholemount retina use, by Coimbra *et al.* 2009, 2012. A computer running the Stereo Investigator software (v2017.01.1 (64-bit), Microbrightfield, USA) coupled to a compound microscope (Zeiss Imager.Z2) equipped with a motorized stage (MAC 6000 System, Microbrightfield, USA) and a digital colour camera (Microbrightfield) was used for the analysis. The contour of each retina wholemount was digitalized using a x5 objective (numerical aperture 0.16) and cells were counted randomly and systematically using a x63 oil immersion objective (numerical aperture 1.40) and the parameters summarised in Tables S1 and S2. The total number of cells for each sample was then estimated by multiplying the sum of the neurons (ganglion cells or photoreceptors) counted by the area of sampling fraction (Coimbra *et al.* 2009; West *et al.* 1991).

The counting frame and grid size were chosen carefully in order to achieve an acceptable Shaeffer’s coefficient of error (CE), while maintaining the highest level of sampling. The CE measures the accuracy of the estimation of the total cell number and it is deemed acceptable below 0.1 (Glaser and Wilson 1998; Slomianka and West 2005). The counting frame was adjusted between life stages to reach an average count of around 40 and 80 cells per sampling site for ganglion cells and photoreceptors respectively, but was kept identical for individuals of the same life stage (Tables S1 and S2, S1). Since fish of similar life stages can have a wide variation of standard lengths, the grid size was adjusted for all individuals to allow sampling of around 200 sites (± 10%) (de Busserolles *et al.* 2014a; de Busserolles *et al.* 2014b).

Three cell types can be found in the ganglion cell layer: ganglion cells, displaced amacrine cells and glial cells. These can usually be distinguished based on cytological criteria (Collin 1988; Collin and Pettigrew 1988c; Hughes 1975) with ganglion cells having an irregular shape, an extensive nucleus, and a larger size compared to smaller, rounder amacrine cells, which have a darker stained appearance, and glial cells having an elongated shape relative to the other two cell types (Fig. 2A). However, since amacrine cells were often difficult to distinguish from ganglion cells in *N. brevirostris*, especially in high density areas, amacrine cells were included in all counts and only glial cells were excluded. The inclusion of amacrine cells in the analysis should not interfere with the overall topography, since the distribution of amacrine cells has been shown to match the ganglion cell distributions in other animals (Coimbra *et al.* 2006; Collin 2008; Collin and Pettigrew 1988c; L. and A. 1987; Bailes *et al.* 2006), and the density of displaced amacrine cells in *N. brevirostris* was relatively low. However, the inclusion of amacrine cells in the ganglion cells counts will contribute to a slight overestimation of ganglion cells densities and ultimately to a slight overestimation of spatial resolving power. For ganglion cell analysis, a sub-sampling was performed in the regions of highest cell density to allow a more accurate estimation of the peak ganglion cell density. The same counting frame parameters as for the whole retina were used for the sub-sampling, but the grid size was reduced by half.

**Fig. 2.**
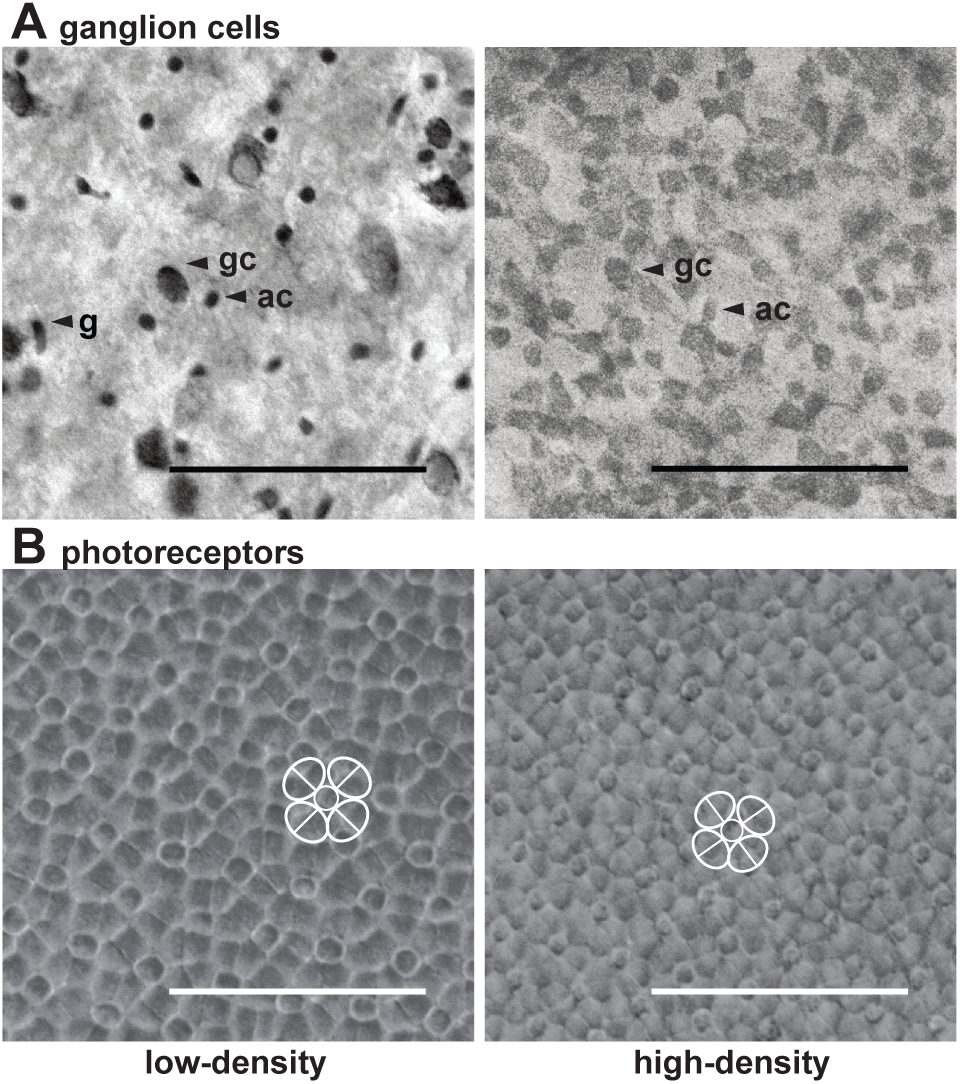
Light micrographs of various retinal layers as found in an adult *N. brevirostris*. (A) Micrographs of the Nissl-stained ganglion cell layer taken in a low-density (nasal part) and a peak-density area (central part) of the retina. Ganglion cells (gc) could clearly be distinguished from glial cells (g) by their round shape and difference in size. Distinguishing amacrine cells (ac) from gc, however, was more difficult. (B) Micrographs of the photoreceptor layer taken in a low-density (nasal part) and a peak-density area (temporal part). Photoreceptors formed a square mosaic with a central single cone (sc) surrounded by four double cones (dc). Scale bar = 50 μm.

Photoreceptor cells, on the other hand, could be distinguished unambiguously into single and double cones (Fig. 2B). Both cone types were counted separately and simultaneously using two different markers to acquire data for single cones alone, double cones alone and total cones (single and double cones).

Topographic maps were created using the statistical program R v.3.4.1 (R Foundation for Statistical Computing, 2012) with the results exported from Stereo Investigator and the R script provided by Garza-Gisholt *et al.* (2014). As for previous retinal topography studies on teleost fishes (de Busserolles *et al.* 2014b; de Busserolles *et al.* 2014a; Dalton *et al.* 2016), the Gaussian Kernel Smoother from the Spatstat package (Baddeley and Turner 2005) was chosen and the sigma value was adjusted to the distance between points (i.e. grid size) for each map (Fig. 3).

**Fig. 3.**
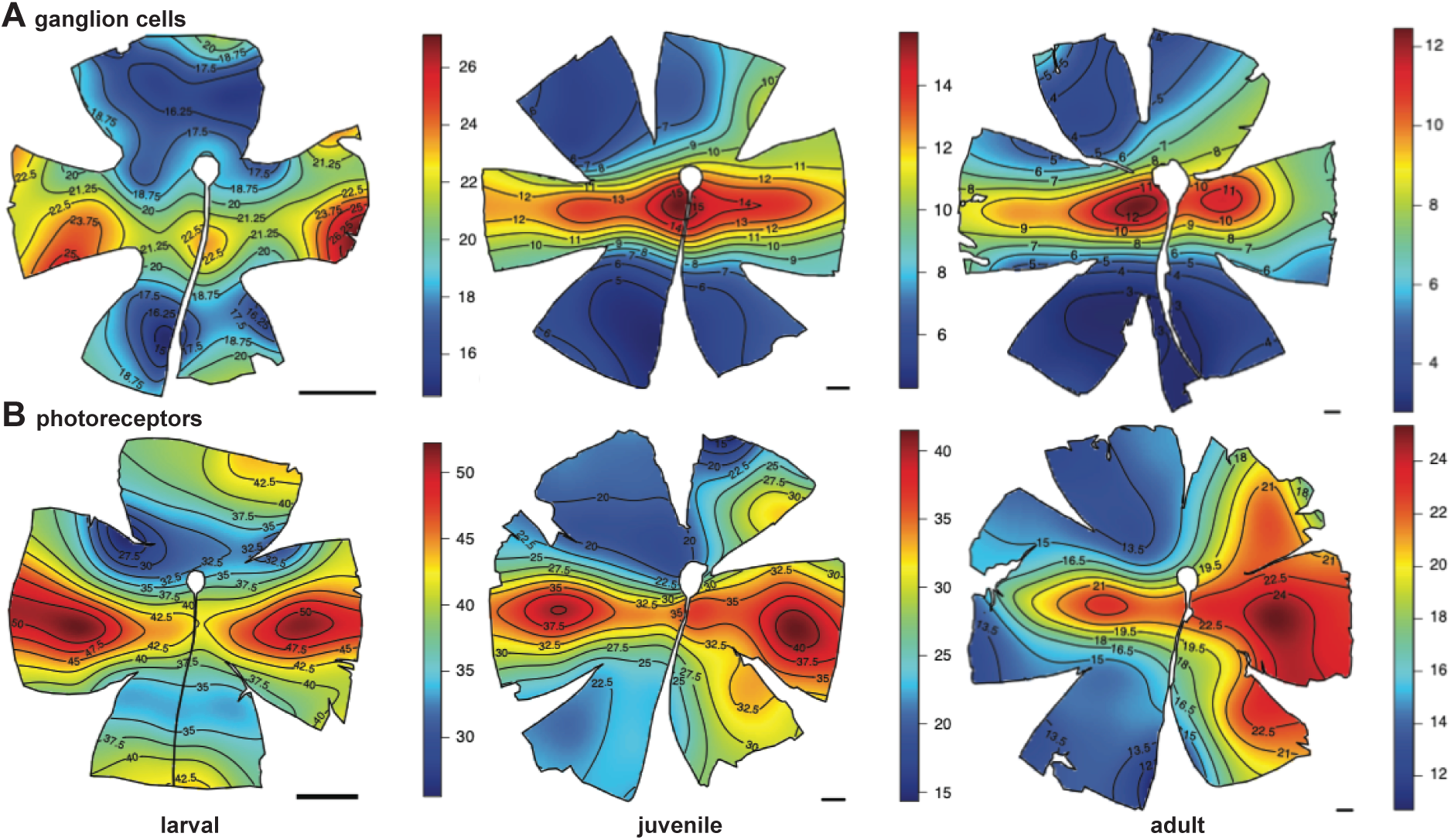
Topographic heat maps of ganglion and photoreceptor-cell distribution. (A) Topographic distribution of retinal ganglion cells revealed a pronounced horizontal streak with a central area of high cell-density in adult and juvenile individuals. The same features, albeit less pronounced, were also present in larvae (see Fig. S1 for maps of additional individuals). (B) Topographic distribution of total photoreceptors (double and single cones) revealed an increase in specialization from two area centralis in larvae to the formation of a horizontal streak and a weak dorsal vertical streak in juveniles. A more pronounced dorsal vertical streak was found in adults (see Figs. S2-S4 for single and double cone maps as well as maps of additional individuals). Black lines represent isodensity contours, and values are expressed in densities x10^3^ cells/mm^2^. V = Ventral, T = temporal. Scale bar = 1 mm.

### Spatial resolving power estimation

The upper limit of the spatial resolving power (SRP) in cycles per degree was estimated for each individual using the ganglion cell peak density as described by Collin and Pettigrew (1989). The following formula was used:

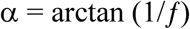

where α is the angle subtending 1 mm on the retina and calculated assuming that f, the focal length of the fish, is 2.55, the standard for teleost fishes according to the Matthiessen ratio (Matthiessen 1882). Then, knowing α, the peak density of ganglion cells (PDG in cells/mm) and the fact that two ganglion cells are needed to distinguish a visual element from its neighbouring element, the SRP in cycles per degree can be calculated as follow:

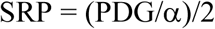

### Transcriptome sequencing, quality filtering, and de-novo assembly

The retinas from different life stages of *N. brevirostris* (adult, n = 3; juvenile, n = 6; larvae, n = 3) were dissected out of the eye cup, total RNA was extracted, and their retinal transcriptomes were sequenced according to Musilova *et al.* (2019). Briefly, total RNA of larval and smaller juvenile retinas was extracted using the RNeasy Mini Kit (Qiagen), and for larger juvenile and adult retinas the RNeasy Midi Kit (Qiagen) according to the manufacturer’s instructions, which included a DNAse treatment. Total RNA concentration and quality were determined using an Eukaryotic Total RNA NanoChip on an Agilent 2100 BioAnalyzer (Agilent Technologies). Juvenile transcriptomes were sequenced in-house at the Queensland Brain Institute’s sequencing facility. For these samples, strand-specific libraries were barcoded and pooled at equimolar ratios and sequenced at PE125 on a HiSeq 2000 using Illumina’s SBS chemistry version 4. Library preparation (strand-specific, 300 bp insert) and transcriptome sequencing (RNAseq HiSeq PE150) for larval and adult individuals was outsourced to Novogene (https://en.novogene.com/).

Retinal transcriptomes were filtered, and *de novo* assembled following the protocol described in (de Busserolles *et al.* 2017). Briefly, raw-reads of transcriptomes were uploaded to the Genomics Virtual Laboratory (GVL 4.0.0) (Afgan *et al.* 2015) on the Galaxy Australia server (https://galaxy-qld.genome.edu.au/galaxy/), filtered by quality using Trimmomatic (Galaxy Version 0.36.4) (Bolger *et al.* 2014) and then *de novo* assembled using Trinity (Galaxy Version 2.4.0.0) (Haas *et al.* 2013).

### Opsin gene mining and phylogenetic reconstruction

Following the protocol in de Busserolles *et al.* (2017), the *N. brevirostris* transcriptomes were mined for their visual opsin genes. Briefly, using the opsin coding sequences from the dusky dottyback, *Pseudochromis fuscus* (Cortesi *et al.* 2016), we searched for the *N. brevirostris* opsin genes by mapping the de-novo assembled transcripts to the *P. fuscus* reference genes using Geneious v.11.1.3 (www.geneious.com). *P. fuscus* was chosen because it is relatively closely related to *N. brevirostris* and because it possesses orthologs from all of the ancestral vertebrate opsin genes (Cortesi *et al.* 2016).

Assemblies based on short-read libraries tend to overlook lowly expressed and similar gene copies and/or short-reads may be misassembled (chimeric sequences); for that reason, a second approach was used to confirm the visual opsin genes of *N. brevirostris.* A manual extraction of the gene copies was performed by mapping raw-reads against the *P. fuscus* references and then moving from single nucleotide polymorphism (SNP) to SNP along the gene taking advantage of paired-end information to bridge gaps between SNPs. The extracted reads were then de-novo assembled and their consensus was used as template against which unassembled reads were re-mapped to elongate the region of interest; this approach eventually lead to a reconstruction of the whole coding region (for details on this approach see de Busserolles *et al.* (2017); Musilova *et al.* (2019)).

Opsin gene identity was then confirmed using BLAST (http://blast.ncbi.nlm.nih.gov/) and by phylogenetic reconstruction to a reference dataset obtained from Genbank (www.ncbi.nlm.nih.gov/genbank/) and Ensembl (www.ensembl.org/) (as per de Busserolles *et al.* (2017)) (Fig. 3). The opsin gene phylogeny was obtained by first aligning all opsin genes i.e. the reference dataset and *N. brevirostris* genes using the L-INS-I settings as part of the Geneious MAFFT plugin v.1.3.7 (Katoh and Standley 2013). jModeltest v.2.1.10 (Darriba *et al.* 2012) was subsequently used to select the most appropriate model of sequence evolution based on the Akaike information criterion. MrBayes v.3.2.6 (Ronquist *et al.* 2012) as part of the CIPRES platform (Miller *et al.* 2010) was then used to infer the phylogenetic relationship between opsin genes using the following parameter settings: GTR+I+G model; two independent MCMC searches with four chains each; 10 million generations per run; 1000 generations sample frequency; and, 25% burn-in.

Opsin gene mining and phylogenetic reconstruction revealed, amongst a number of other visual opsin genes, two *N. brevirostris RH2* paralogs of which one clustered within the *RH2A* clade of other percomorph fishes. However, the phylogenetic placement of the second paralog could not fully be resolved using this approach alone (Fig. 4). Therefore, in order to resolve a more detailed relationship between the two *N. brevirostris RH2* paralogs, we took advantage of the phylogenetic signal within the single exons of the displaced paralog (as per Cortesi *et al.* 2015b; Fig. 5). The five *N. brevirostris RH2-2(B)* exons were obtained by annotating the coding regions of the gene with a *P. fuscus RH2* ortholog. The single exons were separated from one another and inserted as “single genes” in the alignment in a reduced (*RH2* genes only) reference dataset, along with the *N. brevirostris RH2A* gene. The *RH2* specific phylogeny was then reconstructed using MrBayes on the CIPRES platform using the same parameters as before.

**Fig. 4.**
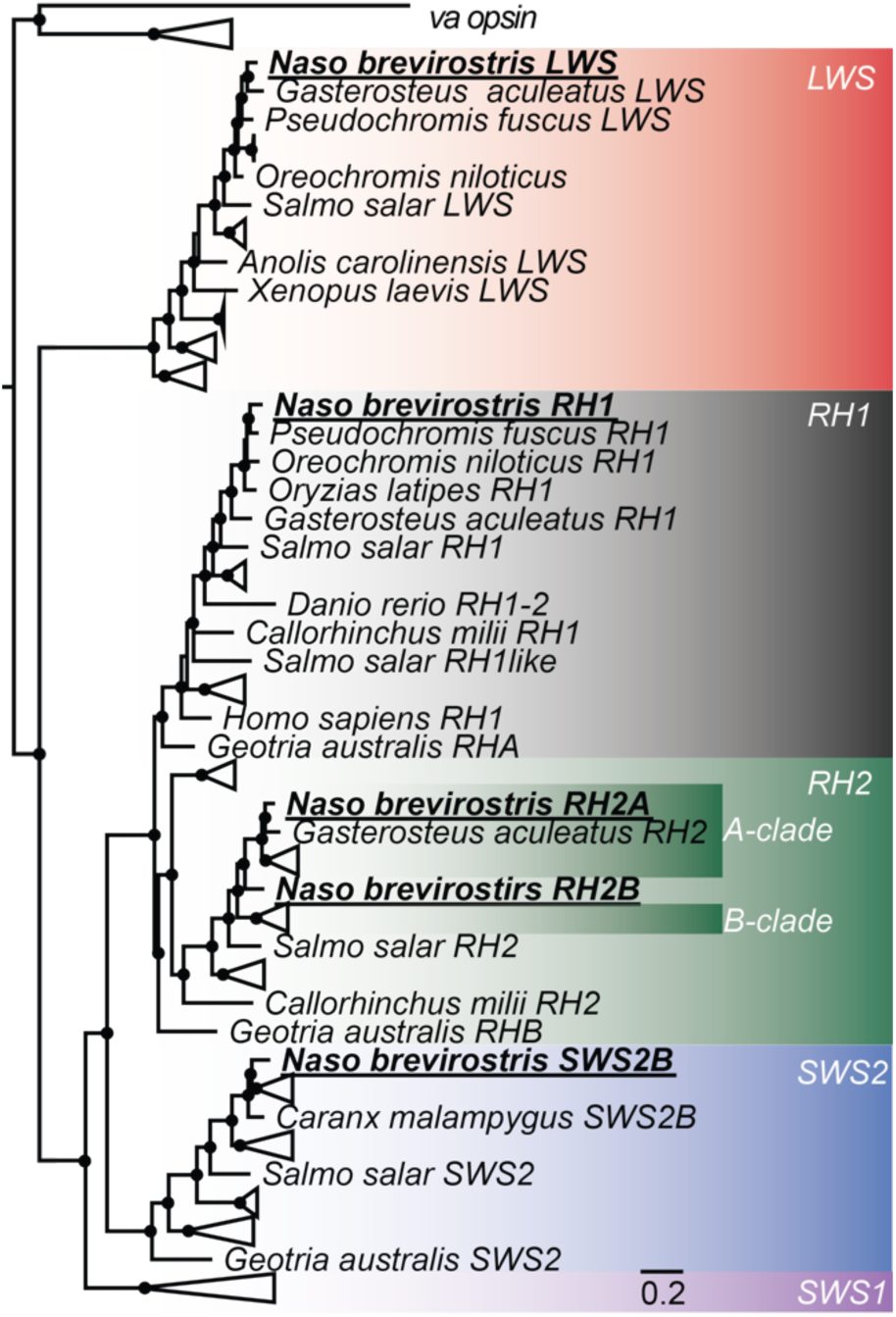
Vertebrate visual opsin gene phylogeny. The different *N. brevirostris* opsin genes which were mined from the retinal transcriptomes are highlighted in bold and belong to four of the five major visual opsin classes. Black spheres indicate Bayesian posterior probabilities > 0.8. Note that the *N. brevirostris RH2B* gene is placed in-between the percomorph *RH2A* and *RH2B* clades (also see Fig. 5). *RH1* = rhodopsin 1 (rod opsin), *RH2* = rhodopsin 2, *SWS2* = short-wavelength-sensitive 2, *LWS* = long-wavelength-sensitive, *va* = vertebrate ancient opsin (outgroup), scalebar = substitution per site. A detailed phylogeny and GenBank accession numbers are shown in Fig. S6.

**Fig. 5.**
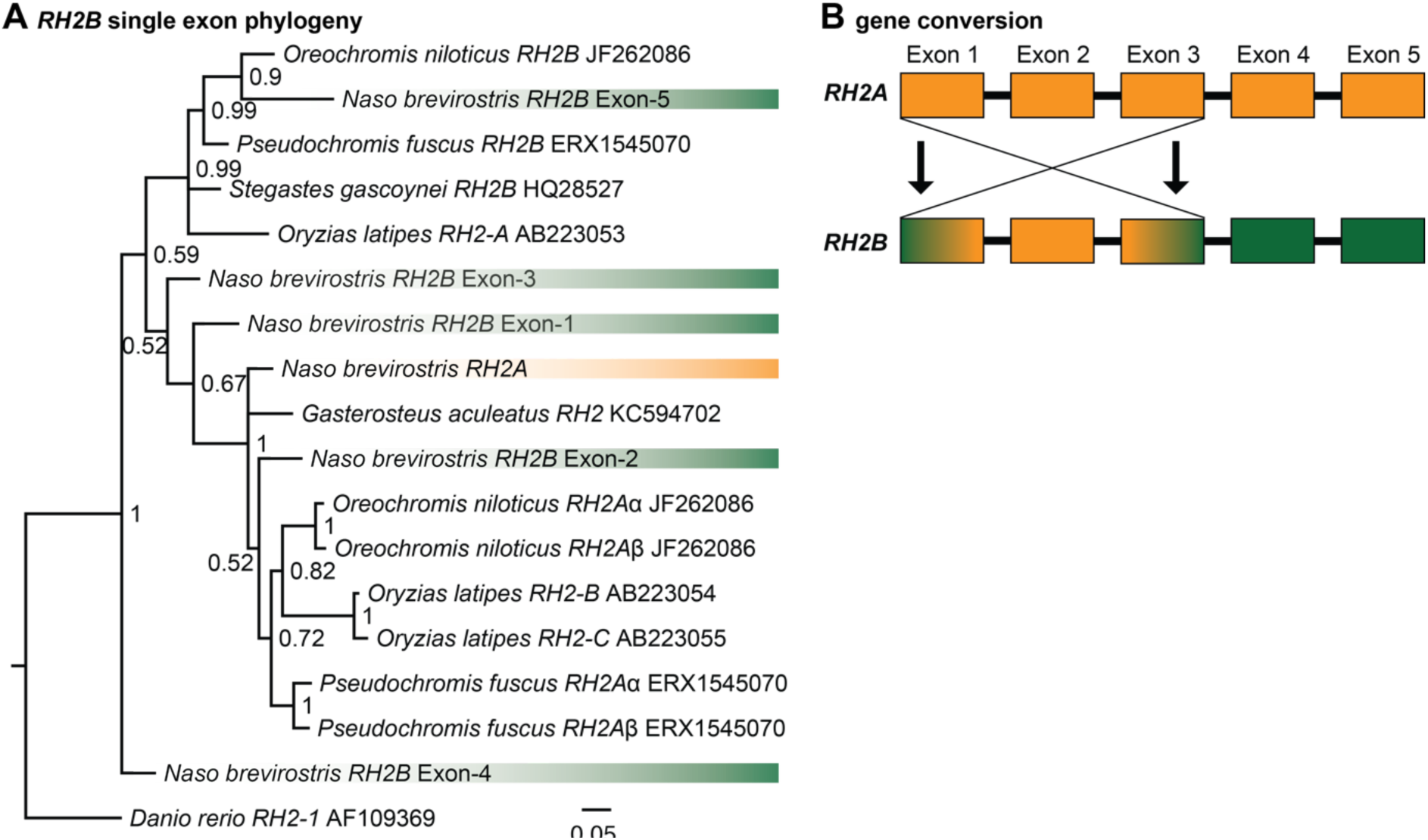
Single-exon green opsin (*RH2*) phylogeny. (A) Using the single exons (green) of the *N. brevirostris RH2B* gene revealed that exons 3-5 cluster within or close to the percomorph *RH2B* clade, whereas exons 1 and 2 cluster close to or within the *RH2A* clade (*N. brevirostris RH2A* in yellow). Note that *Oryzias latipes* genes have a different nomenclature in comparison to the other fish opsin genes. Nodes denote Bayesian posterior probabilities. (B) Illustration of the relationship between the two *RH2* paralogs of *N. brevirostris* based on the single-exon *RH2B* phylogeny. The suggested gene conversion from *RH2A* into Exons 1-3 of *RH2B* makes it near impossible to resolve its phylogenetic position if considering the whole coding region of the gene (also see Fig. 3).

### Opsin gene expression

Quantitative opsin gene expression was assessed by mapping the reads to the assembled coding regions of the *N. brevirostris* opsin genes as per de Busserolles *et al.* (2017). This methodology was used for each individual of the three life stages. Proportional opsin expression for single cones (*p*_i_; SC) and double cones (*p*_i_; DC) for each gene (*i*) was then calculated by first normalizing the number of reads of each gene (R_i_) to the length of each gene specific coding region (cds):

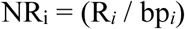

where, R is the number of reads and bp_i_ the number of base pairs in the cds of a gene *i* which was used to normalize the data between the opsins. The proportion of opsin expressed, out of the total normalized expression for single (Tot_SC_) and double cones (Tot_DC_), was then calculated separately. The following formulas were used, depending of which type of cone the gene *i* was expressed in:

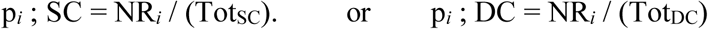

We also calculated the proportional expression of the rod opsin compared to total normalized opsin expression (Tot_Opsin_):

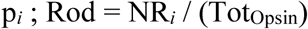

## Results

### Topographic distribution of ganglion cells and spatial resolving power

Topographic maps of ganglion cells (including amacrine cells) for the three life stages of *N. brevirostris* were constructed from Nissl-stained retinal wholemounts. Little variation in topographic distribution of ganglion cells was observed within the same ontogenetic stage. Therefore, only the topographic map of one individual per life stage is presented here (Fig. 3A), and the results for the remaining individuals are shown in Supplementary Fig. S1.

Differences in retinal topography were mainly found between the larval stage and the two later stages (Fig. 3A). In general, the larval retina showed less specializations compared to juvenile and adult retinas. In the larval retina, an onset of a horizontal streak was observed with the highest cell density found in the central meridian of the retina (1.5x increase compared to the areas with the lowest cell densities). Within this weak streak, three areas of high cell densities were found; in the nasal, central and temporal parts of the retina. However, these areas of high cell densities are to be taken with caution due to the limitations of the Nissl-staining protocol for very small retinas. Larval retinas were challenging to prepare and analyse due to their small size and thus, the higher amount of shrinkage present after staining. After several attempts with different larvae, only one larval retina was deemed acceptable for analysis. Even for this individual, the areas of high cell densities in the nasal and temporal part of the retina are questionable, since they are very close to the retinal borders and therefore could be the result of shrinkage. A prominent horizontal streak along with a centralized area centralis (the area centralis had a 2.5-3x increase in cell density compared to the areas with the lowest cell densities) was present in the juvenile and adult individuals. Similarly to the larvae, the streak in juveniles and adults was located on the central meridian of the retina extending to the nasal and temporal margins. Although slightly different patterns were found for each life stage, they all showed a higher ganglion cell density in the central area close to the optic nerve, accompanied by a horizontal streak (Fig. 3A).

The total number of ganglion cells increased with the size of fish and ranged from 208,975 cells for the larval individual, over ~1,600,000 cells for juveniles, to ~2,100,000 cells for adults (Table 1). Conversely, the mean cell density decreased with the size of the fish ranging from 19,439 cells/mm^2^ in the larval individual, over ~8,500 cells/mm^2^ in juveniles, to ~5,000 in cells/mm^2^ in adults. Peak cell density also decreased through development, from 30,400 cells/mm^2^ in the larval individual, to ~23,000 cells/mm^2^ in juveniles, and ~ 20,500 cells/mm^2^ in adults.

**Table 1.** Summary of ganglion cell quantitative data obtained using the optical fractionator method on the wholemounted retinas of three developmental stages of *N. brevirostris*.

| Stage | Individual | Peak cell density,<br>(cells/mm<sup>2</sup>) | Mean cell density<br>(cells/mm<sup>2</sup>) | Total cells | Lens Ø<br>(mm) | SRP |
| --- | --- | --- | --- | --- | --- | --- |
| Adult | ID1 | 20,617 | 5,340 | 2,034,000 | 6.6 | 10.6 |
|  | ID2 | 20,370 | 4,772 | 2,100,825 | 6.9 | 11.0 |
| Juvenile | ID1 | 23,750 | 8,130 | 1,617,968 | 4.7 | 8.1 |
|  | ID2 | 24,531 | 8,688 | 1,737,656 | 4.9 | 8.6 |
|  | ID3 | 21,875 | 8,584 | 1,450,625 | 4.5 | 7.5 |
| Larvae | ID1 | 30,400 | 19,439 | 208,975 | 1.4 | 2.98 |

SRP = spatial resolving power, Ø = diameter

Based on the peak of ganglion cells densities, the SRP of *N. brevirostris* ranged from 2.98 cycles per degree in the larval individual, over ~8.0 cycles per degree in juveniles, to a maximum of 11.0 cycles per degree in adults (Table 1). Overall, SRP or visual acuity in *N. brevirostris* increased with the size of the fish with very little variation found within ontogenetic stages (Fig. S2).

### Topographic distribution of cone photoreceptors

The density and topographic distribution of cone photoreceptors (double and single cones), was assessed in the three life stages of *N. brevirostris*. Double and single cones were arranged in a square mosaic, with one single cone at the centre of four double cones (Fig. 2B). This pattern was consistent throughout the entire retina, thus providing a ratio of double cones to single cones of 2:1. As a consequence of this regular arrangement, the topographic distribution of single cones, double cones and total cones was identical. Moreover, similar to the ganglion cell topography, little variation in topographic distribution of cone photoreceptors was observed within the same ontogenetic stage. Therefore, only the total cone topographic map of one individual per life stages is presented here (Fig. 3B). The remaining maps (i.e., for single and double cones separately, and maps of all individuals) are provided in the Supplementary Figs. S3 – S5.

The topographic distribution of cone photoreceptors varied between stages with a pronounced increase in specialization from the larval to the juvenile stage and smaller changes thereafter (Fig. 3B). Larvae had a weak horizontal streak in the central meridian as well as two area centralis, one in the nasal part and one in the temporal part of the retina. One of the two analysed larval individuals also showed a dorsal increase in cell density (Fig. S3f). However, this apparent increase in cell density was likely caused by an artefact from not properly flattening the dorsal part of the retina during mounting and should therefore be disregarded (Figs. S3 – S5). Compared to the larvae, juveniles had a more pronounced horizontal streak in the central meridian. The two area centralis were still present but the nasal one was less pronounced, and the peak cell density was found in the temporal area centralis. Moreover, a weak vertical streak could be seen in the temporal part of the retina, extending from the dorso-temporal area to the ventral-temporal area. In adults, the horizontal streak in the central meridian was still present but did not extend as far into the nasal part as in the juveniles. Moreover, the vertical streak was more prominent compared to the one found in juveniles resulting in a large area of high cell density in the temporal region (Fig. 3B). The continuous nature of the transition between juvenile and adult specializations is highlighted by the topography of individuals of intermediate sizes (Figs. S3-S5). For example, the horizontal streak was less pronounced in the nasal part of a larger (Fig. S3c) compared to a smaller juvenile (Fig. S3d). Conversely, the vertical streak in a smaller adult (Fig. S3b) was still developing compared to the one found in a larger adult (Fig. S3a).

Similar to the ganglion cells (Table 1), the total number of photoreceptors increased with the size of the fish ranging from ~650,000 cells in larvae, over ~4,300,000 cells in juveniles, to ~5,700,000 cells in adults (Table 2). A large difference in the total number of photoreceptors was found between the two juvenile individuals. This difference is likely due to the size difference between these individuals. Photoreceptor peak cell densities decreased with the size of the fish, ranging from ~69,000 cells/mm^2^ in larvae, over ~51,000 cells/mm^2^ in juveniles, to ~34,000 in cells/mm^2^ in adults (Table 2).

**Table 2.** Summary of photoreceptor quantitative data obtained using the optical fractionator method on the wholemounted retinas of three developmental stages of *N. brevirostris*.

| Stage | Individual | Total DC | Peak DC<br>(cells/mm<sup>2</sup>) | Total SC | Peak SC<br>(cells /mm<sup>2</sup>) | Total cones | Peak TC<br>(cells mm<sup>2</sup>) |
| --- | --- | --- | --- | --- | --- | --- | --- |
| Adult | ID3 | 3,738,296 | 23,703 | 1,926,513 | 12,098 | 5,664,809 | 35,801 |
|  | ID4 | 3,820,839 | 21,208 | 1,972,975 | 12,345 | 5,793,814 | 33,553 |
| Juvenile | ID3 | 2,466,222 | 34,218 | 1,283,524 | 18,125 | 3,749,746 | 52,343 |
|  | ID4 | 3,212,099 | 32,968 | 1,667,475 | 17,187 | 4,879,574 | 50,155 |
| Larvae | ID2 | 452,010 | 45,432 | 231,371 | 23,703 | 683,381 | 69,135 |
|  | ID3 | 413,476 | 45,432 | 211,105 | 23,704 | 624,581 | 69,136 |

DC = double cones; SC = single cones

The total number of cone photoreceptors was greater compared to the total number of ganglion cells, indicating a high summation ratio between the two cell types. For one individual (larva ID3), the distribution of both ganglion cells and photoreceptors were analysed, which allowed to estimate the summation ratio between photoreceptors and ganglion cells in low- and high-density areas, respectively. For this individual, the summation ratio was found to be as low as 2.3 in the central part and as high as 5.4 in the ventral-temporal part of the retina.

### Visual opsin genes and their expression in Naso brevirostris

*N. brevirostris* were found to mainly express four opsin genes in their retinas. Independent of ontogeny, these were the ‘blue-violet’ *SWS2B*, the ‘greens’ *RH2B* & *RH2A*, and the rod opsin *RH1*. The ‘red’ *LWS* was also found to be expressed, albeit at very low levels in all stages (0.1 – 6.5 % of total double cone opsin expression; Fig. 6A, Table S3). The phylogenetic reconstruction based on the full coding regions of the genes confirmed the positioning of all genes within their respective opsin class (Fig. 4). However, for *RH2B* in particular the resolution between *RH2* specific clades was poor (Fig. 4, Fig. S6). This was resolved using the exon-based approach which showed the placement of some of the *N. brevirostris RH2B* exons within a greater percomorph *RH2B* clade (Fig. 5A). Moreover, we found evidence for substantial gene conversion affecting this gene with the placement of Exons 1 and 2 close to, or within, the *RH2A* clade (Fig. 5B).

**Fig. 6.**
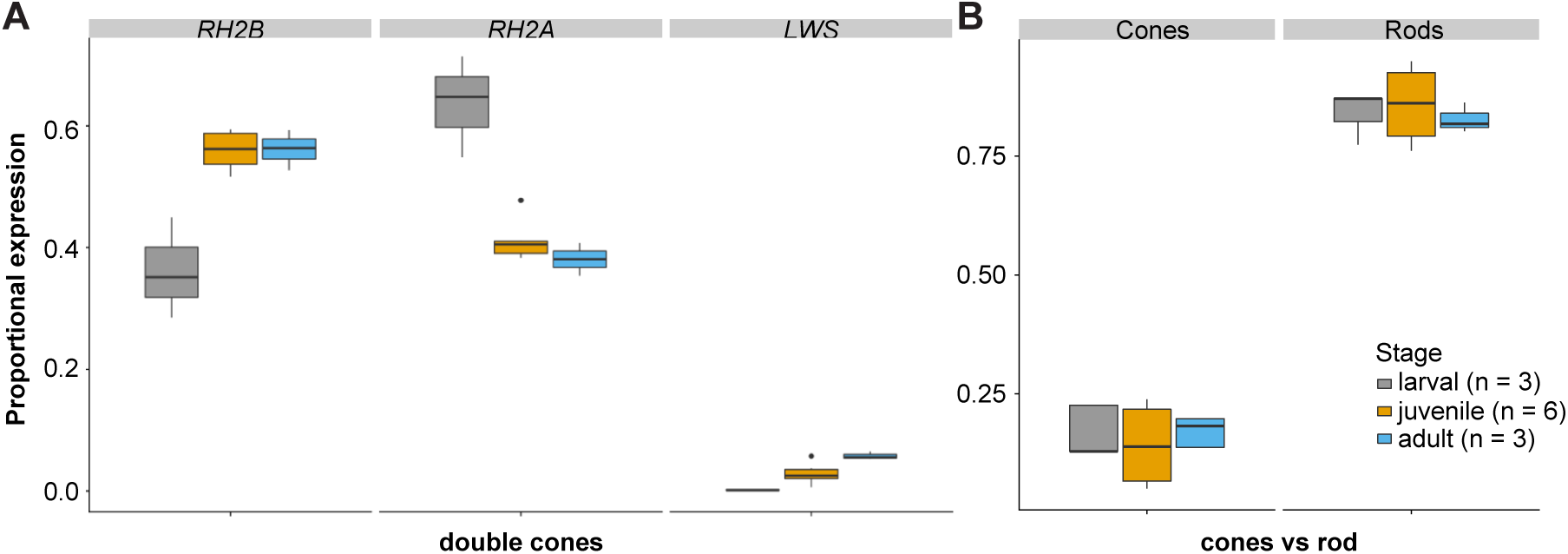
Proportional expression of *N. brevirostris* opsin genes. (A) Independent of ontogenetic stage, *N. brevirostris* expressed the *SWS2B* single cone (100% of single cone expression; see Table S3 for details) and three double cone opsin genes: *RH2B*, *RH2A*, and *LWS*. The proportional expression of double cone opsins revealed a change in expression of the two *RH2* genes between the larval and the juvenile stage as well as a steady increase in the expression of *LWS* with development. (B) The proportional expression of cone (*SWS2B*, *RH2B*, *RH2A*, *LWS*) versus rod opsin (*RH1*) remained similar throughout ontogeny. The box indicates Q2 and Q3, with the line indicating the median and the whiskers indicating Q1 and Q4 of the data.

Quantitative opsin gene expression revealed that *SWS2B* was the only single cone gene and thus, expressed at 100% in all developmental stages (Table S3). Of the double cone opsins, there was a change in expression for the *RH2* genes with ontogeny. During the larval stage (n = 3), *RH2B* (mean ± s.e., 36.2 ± 4.8%) was less highly expressed compared to *RH2A* (63.6 ± 4.8%). The opposite pattern was found in the juvenile (n = 6) and adult stages (n = 3), where *RH2B* was the highest expressed of all double cone opsins genes (juvenile: 56 ± 1.3%; adult: 56.1 ± 1.9%). *RH2A* in the juvenile (41.2 ± 1.4%) and adult (38.1 ± 1.6%) stages was less highly expressed. Despite *LWS* being lowly expressed in all stages, there was a noticeable increase in expression with development (larval: 0.2 ± 0.0%; juvenile: 2.8 ± 0.7%; adult 5.8 ± 0.4%; Fig. 6A). Rod opsin (*RH1*) expression was substantially higher compared to the cone opsin expression in all stages (82 – 86% for all stages) (Fig. 6B).

## Discussion

The visual systems of fishes often change through development when transitioning from one habitat to another. These changes are usually associated with a shift in light environment e.g., when moving from the open ocean to a coral reef, but possibly also with changes in diet and predation pressure (Sale 2013). Our objective was to assess the visual system development in the spotted unicornfish, *N. brevirostris*. *N. brevirostris* experiences multiple changes in habitat, diet and morphology throughout ontogeny (from larval to adult stages; Fig. 1) making it a prime candidate to study visual system changes on the reef.

### Ganglion cell topography

Retinal topography is an effective method to identify visual specializations and recognise the area of the visual field a species is most interested in (Hughes 1977; Collin and Pettigrew 1988a; Collin 2008). In marine fishes, visual specializations have been found to correlate with the structure and symmetry of the environment they live in and/or with their feeding behaviour (Collin and Pettigrew 1988a, 1988b; Ito and Murakami 1984; Shand 1997; Caves *et al.* 2017). In this study we show that the *N. brevirostris* eye possesses a horizontal streak in all life stages (Fig. 3A). This type of specialization has previously been found in species living in open environments where an uninterrupted view of the horizon, defined by the sand-water or air-water interface, is present (Collin and Pettigrew 1988b). Since *N. brevirostris* spends much of its life (larval and adult stages) searching for prey in the water column, having a pronounced horizontal streak is likely to increase feeding and predator surveillance capabilities by allowing it to scan the horizon without using excessive eye movements (Collin and Shand 2003). Moreover, at the larval stage this type of specialization may also enable fish to scan the environment when searching for a reef habitat to settle on. On the contrary, a horizontal streak does not seem to match the visual needs at the juvenile stage during which *N. brevirostris* lives in close association with the reef i.e., in a more enclosed 3D environment. At this life stage, we would have expected to find one (or multiple) area centralis and no horizontal streak; a common feature in fishes that live close to, or within the reef matrix (Collin and Pettigrew 1988a; Collin and Pettigrew 1988c). Compared to the lifespan of these fishes (up to 20 years), the juvenile stage is relatively short (~ 3 years; Choat and Axe 1996), which may explain the maintenance of the horizontal streak throughout development.

On top of having a well-defined streak, the ganglion cells in the juvenile and adult stages also formed an area centralis in the central part of the retina (Fig. 3A). This is very unusual, as in coral reef fishes an area centralis is normally found in the temporal zone. Such a temporal area centralis receives information from the nasal visual field, and thus is usually correlated with feeding and predator avoidance in front of the fish (Collin and Pettigrew 1988a; Collin and Pettigrew 1989; Fritsch *et al.* 2017; Fritsches *et al.* 2003; Shand *et al.* 2000). The type of specialization found in *N. brevirostris* seems to be correlated with its unusual visual behaviour as fishes are found to examine objects side-on (V.T. pers. observation). A possible explanation for this peculiar behaviour is that due to its protruded snout, which grows through development, the frontal image might be partially blocked and stereoscopic vision may be impaired or even impossible (Purcell and Bellwood 1993). Although the visual field of *N. brevirostris* was not investigated in this study, Brandl and Bellwood 2013 suggested that the protruded snout found in many *Naso* species indeed prevents an overlap of their horizontal field of view. Similar to the monocular vision found in hammerhead sharks (McComb *et al.* 2009; Lisney and Collin 2008), increasing visual acuity in the central part of the retina would thus maximise a sideward oriented visual field. Together with a pronounced visual streak, these two specializations are likely to enable *N. brevirostris* to accurately navigate both within the complexity of the reef as well as in open water.

### Photoreceptor topography

Similar to the ganglion cell topography, the photoreceptor topography also varied mostly between the larval and subsequent stages (Fig. 3B). Larval fishes had two well defined area centralis in the nasal and temporal zones, which comply with two of their main ecological needs: feeding (temporal; looking forward) and predator avoidance (nasal; looking backwards) (Collin and Pettigrew 1988a; Fortier and Harris 1989; Boehlert 1996). These high-density regions were not matched by the ganglion cell topography and as such, are likely to provide areas of higher sensitivity (i.e., areas of high photoreceptor to ganglion cell ratio; Walls 1942). Moreover, although larval fishes rely mainly on olfactory cues to zoom in on a suitable habitat for settlement (Lecchini *et al.* 2005b), the temporal area centralis in particular might also assist when searching for said habitat over longer distances (Mouritsen *et al.* 2013).

The two area centralis were no longer present in bigger fishes, but instead, at the juvenile and adult stage, *N. brevirostris* showed a pronounced horizontal streak (Fig. 3B). Additionally, a temporal vertical specialization became apparent at the juvenile stage and more pronounced in adults. Such a double streak specialization, with a vertical and a horizontal component, is a first in coral reef fishes. *N. brevirostris* adults live on the coral reef slope/wall, and move up and down the wall (from 2 - 122 m) while foraging and searching for mates (Mundy 2005). As such, in line with the terrain hypothesis (Hughes 1977), the evolution of this vertical specialization is likely a result of the vertical component in their visual environment.

A difference in the topography of ganglion cells and photoceptors means that the summation ration between the cell types i.e., the sensitivity and spatial resolution of the retina, also differs depending on the visual field in question. For example, high photoreceptor densities and comparable low ganglion cell densities in the ventral-temporal and dorsal-temporal parts of the vertical streak confer higher sensitivity to these two areas (Walls 1942). Theoretically, this enables juvenile and adult *N. brevirostris* to detect even small differences in luminance, which may help to detect well camouflaged predators against the reef wall. A high density of both photoreceptor and ganglion cells found in the centre of the retina, on the other hand, confers a low summation ratio which leads to an increase in visual acuity (Walls 1942). This area of high acuity may help fish to identify conspecifics and also to distinguish between food items (Cronin *et al.* 2014).

To summarize, the photoreceptor topographies of *N. brevirostris* may be adapted to the habitat in both the larval and the adult stage. Juveniles live in a more enclosed, 3D coral reef environment compared to the other two life stages. Therefore, we would have expected the juvenile visual system to reflect its habitat by having a less developed streak and a more pronounced area centralis. Similar to the ganglion cell topography, the lack of a distinct area centralis in the retina may be explained by the juvenile stage only lasting a fraction of the lifespan of *N. brevirostris* (Choat and Axe 1996). The relatively short period of time spent in a habitat rich in shelter and food enables the fish to grow big enough to avoid most predators (Lasiak 1986; Barnes and H ghes 1999). During this time, juvenile *N. brevirostris* mostly feed on benthic algae, which do not require a highly specialized visual system in terms of retinal topography (Randall *et al.* 1997; Collin and Pettigrew 1988b, 1988a; Caves *et al.* 2017). A such, instead of changing the visual system multiple times, it is likely more energy efficient to maintain (or slightly adjust) a visual system that is optimally adapted for both the larval and adult stages.

### Visual acuity

The visual acuity of *N. brevirostris* was found to increase through development (Table 1). This seems to be a common feature in coral reef fishes, as a higher acuity often correlates with an increase in eye size during growth. The benefit of having a higher visual acuity is that, as fishes grow and expand their home ranges, it increases the distance at which visual objects such as predators, conspecifics, and food can be detected (Shand 1997; Caves *et al.* 2017). Accordingly, like in other coral reef fish larvae (Shand 1997), the acuity of *N. brevirostris* larvae was relatively poor (2.98 cycles per degree). The overabundance of zooplankton in their habitat means that larval coral reef fishes can wander instead of using a lock-and-pursuit feeding behaviour i.e., they do not need to spot their food from a distance, but rather bump into it while floating in the plankton (Fortier and Harris 1989; Evans and Fernald 1990). Once settled on the reef, the visual acuity of *N. brevirostris* starts to increase in line with their growth (Fig. S5). Adult *N. brevirostris* were found to have a similar visual acuity (~11 cycles per degree) to other reef fishes of that size such as in the clown triggerfish, *Balistoides conspicillus*; a species that also inhabits the reef slope and shows a pronounced horizontal streak (Collin and Pettigrew 1989).

### Opsin gene evolution

Phylogenetic reconstruction showed that the *N. brevirostris* visual opsins belong to the opsin gene clades usually found within percomorph fishes (Fig. 4). However, within the *RH2* genes, an exon-based phylogeny revealed that the *N. brevirostris RH2B* gene is likely to have undergone substantial gene conversion (Fig. 5). As such, it occurs that parts of its first and second exon have been acquired from the *RH2A* paralog explaining its phylogenetic uncertainty when using whole coding region-based reconstructions (Fig. 4, Fig. S6). This is not that surprising, since *RH2* duplicates in teleosts are commonly found in tandem (e.g., Musilova *et al.* 2019) and, as is the case for other teleost opsin genes (Cortesi *et al.* 2015b; Hofmann and Carleton 2009), frequently experience gene conversion (Cortesi *et al.* 2015b; Escobar-Camacho *et al.* 2016; Hofmann *et al.* 2012). This phenomenon is thought to be one of the main mechanism for concerted evolution in small gene families which often originate from tandem duplications (Ohta 1983; Li 1997) and could help to preserve gene function by repairing null-mutations (Innan 2009) or by resurrecting previously pseudogenized gene copies (Cortesi *et al.* 2015b). Since the *RH2* opsin genes are highly expressed in *N. brevirostris* they seem rather important for their visual ecology, and it is therefore likely that gene conversion played a major evolutionary role in maintaining their function.

### Heterochrony of opsin gene expression

Based on opsin gene expression, *N. brevirostris* could be behaviourally trichromatic (i.e. has three spectral sensitivities) for all three developmental stages, with the ‘violet’ *SWS2B*, and the ‘blue-green’ *RH2B* and *RH2A* genes being expressed in sufficient quantity to enable this level of chromatic analysis. Supporting these findings, microspectrophotometry (MSP) in adults of two closely related *Naso* species (*N. literatus* and *N. unicornis*; Sorenson *et al.* 2013) found three cone photoreceptors with spectral sensitivities ~ 420 nm λ_max_ for single cones, and ~ 490 nm λ_max_ and ~ 515 nm λ_max_ for the accessory and principle members of double cones, respectively (Losey *et al.* 2003). A short-shifted visual system with high sensitivtiy in the violet to green range might benefit feeding on zooplankton and gelatinous prey during the larval and adult stages of *N. brevirostris* (Marshall *et al.* 2003). However, it seems at odds with the mainly algivorous diet of the juvenile stage, where a red-shifted visual system would be of advantage (Stieb *et al.* 2017; Cortesi *et al.* 2018).

Ontogenetic studies on opsin gene expression in African cichlids highlighted three main developmental patterns: i) a ‘normal’ development with a display of different gene sets in the larval, juvenile and adult stages; ii) a neotenic development in which the fish retains the larval opsin gene expression throughout its life, or slowly progresses to a slightly different juvenile opsin set; and, iii) a direct development with the fish expressing the adult gene set already at the larval stage (Carleton *et al.* 2008; O’Quin *et al.* 2011). Neotenic and direct development are forms of heterochrony (O’Quin *et al.* 2011). In *N. brevirostris*, we would have expected scenario i), with different opsin sets expressed at different life stages as a consequence of being exposed to varying environments and feeding habits through development. Conversely, we found evidence for a neotenic development (scenario ii), with a slight shift in opsin gene expression between the larval and the juvenile stages (decrease in *RH2A* and increase in *RH2B* expression), which was then retained through to the adult stage (Fig. 5). Neoteny in opsin gene expression was also found in some cichlids from Lake Malawi (Carleton and Kocher 2001; Carleton *et al.* 2008). Similar to the light environment found on the reef (Marshall *et al.* 2003), these fishes inhabit clear water lakes throughout their life (Carleton *et al.* 2008). The consistency in photic habitat as well as zooplanktivory are thought to drive the neotenic development in these cichlids (Carleton *et al.* 2008). Likewise, feeding on zooplankton during larval and adult stages as well as little changes in light habitat post settlement might be responsible for the neotenic expression patterns found in *N. brevirostris*. Supporting the molecular findings, the retinal topography, and especially the ganglion cell topography, also showed a neotenic development, changing slightly from the larval to the juvenile stage with no major changes thereafter (Fig. 3).

A shift in the expression of *RH2B* and *RH2A*, as seen between the larval and later *N. brevirostris* stages, can also be found in coral reef damselfishes (Pomacentridae) (Stieb *et al.* 2016). On shallow, clear coral reefs a broad spectrum of light is available (Marshall *et al.* 2003). However, with increasing depth the long and short ends of the spectrum are cut off due to absorption and scattering through interfering particles, resulting in a blue mid-wavelength saturated light environment (Smith and Baker 1981). Consequently, in an attempt to maximise photon catch, some damselfish species were found to increase the expression of the blue-sensitive *RH2B* gene and simultaneously decrease the expression of the green-sensitive *RH2A* gene with increasing depth (Stieb *et al.* 2016). In *N. brevirostris*, the shift in expression of *RH2* genes occurs between the larval and juvenile stages where depth differences do not seem that relevant. In lieu of depth, individuals migrate from a pelagic blue-shifted open water environment to the more green-shifted light environment of the coral reef (Marshall *et al.* 2003). This could in theory explain the high *RH2A* expression in larval fish at the settlement stage, however, it does not explain the increase in *RH2B* expression post settlement (Fig. 6A). An increasing number of fishes are found to change their opsin gene expression to tune photoreceptors to the prevailing photic environment (e.g., Fuller *et al.* 2004; Hofmann *et al.* 2010; Nandamuri *et al.* 2017; Shand *et al.* 2008; Stieb *et al.* 2016; Luehrmann *et al.* 2018; Härer *et al.* 2017). At the opposite end of the spectrum, opsin gene expression might be pre-programmed either by phylogeny or on a species by species basis, as exemplified by only some damselfishes changing expression with depth (Stieb *et al.* 2016). It is therefore possible that opsin gene expression in *N. brevirostris* is under phylogenetic control and that changes in photic environment contribute very little to opsin gene expression in this case.

*N. brevirostris* was not found to express the UV-sensitive *SWS1* gene at any of the developmental stages. *SWS1* expression is often found in larval fishes and more generally in fishes feeding on zooplankton, with UV-vision thought to increase the detectability of this food source (Sabbah *et al.* 2010; Novales-Flamarique and Hawryshyn 1994). Since *N. brevirostris* feeds on zooplankton at both larval and adult stages (Choat *et al.* 2002; Choat *et al.* 2004), the lack of *SWS1* expression seems striking. However, it does support ocular media measurements which revealed UV-blocking lenses in both larval and adult *N. brevirostris* (Siebeck and Marshall 2007). UV-blocking lenses seem common in many bigger coral reef fishes, which is thought to enhance sighting distance by reducing chromatic aberration and scatter, as well as protecting the eye from the damage caused by these high intensity wavelengths (Siebeck and Marshall 2001). Instead, the expression of the violet sensitive *SWS2B* gene, since its spectral absorption curve reaches into the near-UV (Losey *et al.* 2003), may be sufficient to increase the discrimination of zooplankton from the water background while foraging.

We furthermore found low expression of the red-sensitive *LWS* gene (<6%) at all developmental stages. This suggests that *LWS* expression is either restricted to certain areas of the retina, interspersed at low frequency across the retina, or some photoreceptors might co-express *LWS* with an *RH2* gene (e.g., Dalton *et al.* 2014; Cortesi *et al.* 2016; Torres-Dowdall *et al.* 2017). MSP in related *Naso* species did not show any long-wavelength-sensitive photoreceptors, nor did it show any evidence for opsin co-expression (i.e., red-shifted unusually broad absorbance peaks) (Losey *et al.* 2003). Since this technique only samples as subset of the photoreceptors across the retina, it might be that the photoreceptors containing this pigment were missed due to their low number or that *LWS* was simply not expressed in these fishes.

It is possible that the *LWS* expression found here is just a by-product of the way opsin gene expression is controlled and that it does not serve any ecological function. Nevertheless, *LWS* expression did increase with development. Hence, an alternative explanation might be that *LWS* is co-expressed with an *RH2* gene, which has been shown to increase achromatic discrimination in cichlids (Dalton *et al.* 2014). Moreover, *LWS* expression has recently been shown to be correlated to algal feeding in damselfishes (Stieb *et al*. 2017), and blennies (Cortesi *et al.* 2018). Since *N. brevirostris* juveniles feed on algal turf, a slight increase in *LWS* expression at this stage, may improve feeding efficiency due to the increased contrast of algae against the reef background (Stieb *et al.* 2017; Marshall *et al.* 2003; Cortesi *et al.* 2015a). In situ hybridisation studies (e.g., Dalton *et al.* 2014, 2016; Torres-Dowdall *et al.* 2017; Stieb *et al.* 2019) coupled with behavioural colour-vision experiments (e.g., Cheney *et al.* 2019) will be needed in the future to assess the distribution and function of the various opsin genes and ultimately the colour vision system of *N. brevirostris*.

## Conclusion

Using a multidisciplinary approach, we analysed the ontogeny of the visual system of *Naso brevirostris*. Minor ontogenetic changes in retinal topographies and opsin gene expression were only found after the larval stage, which did not match the initial hypothesis of an adaptation to each developmental stage. Therefore, both retinal topography and opsin expression undergo a neotenic development already possessing the adult, ‘final’ visual system early on in development. This is contrary to what was found in other reef fishes (Shand *et al.* 2008; Suresh and Julia 2001; Cortesi *et al.* 2016) and highlights the need for a comprehensive analysis of visual ontogeny across the reef fish community.

## Acknowledgements

We would like to thank Sara Stieb, Vivian Rothenberger, Simon Dunn, and the staff of Tetiaroa Society (Moana LeRohellec) and of CRIOBE (Camille Gache) for assistance with specimen collection. We furthermore thank the staff at the Lizard Island Research Station for logistical support, and Janette Edson from the Queensland Brain Institute’s (QBI) sequencing facility for library preparation and RNA sequencing. We also acknowledge QBI’s Advanced Microscopy Facility for the use of the Stereo Investigator (software v. 2017.01.1), generously supported by the Australian Government through the ARC LIEF grant LE100100074.

## Author Contributions

F. C. conceived the study and designed the experiments together with F.d.B and N.J.M. V.T., F.d.B and F.C. performed the experiments and analysed the data. All authors contributed to specimen collection. V.T. wrote the initial draft of the manuscript and all authors agreed to the final version of the manuscript.

## Data Accessibility

Raw-read transcriptomes (PRJ tba) and single gene sequences (#tba) are available through GenBank (https://www.ncbi.nlm.nih.gov/genbank/). Gene alignments and single gene phylogenies can be accessed through Dryad (#tba). All other data is given either in the main manuscript or the supplementary material.

